# Toden-E: Topology-Based and Density-Based Ensembled Clustering for the Development of Super-PAG in Functional Genomics using PAG Network and LLM

**DOI:** 10.1101/2024.10.20.619308

**Authors:** Qi Li, Cody Nichols, Robert S Welner, Jake Y. Chen, Wei-Shinn Ku, Zongliang Yue

## Abstract

The integrative analysis of gene sets, networks, and pathways is pivotal for deciphering omics data in translational biomedical research. To significantly increase gene coverage and enhance the utility of pathways, annotated gene lists, and gene signatures from diverse sources, we introduced pathways, annotated gene lists, and gene signatures (PAGs) enriched with metadata to represent biological functions. Furthermore, we established PAG-PAG networks by leveraging gene member similarity and gene regulations. However, in practice, high similarity in functional descriptions or gene membership often leads to redundant PAGs, hindering the interpretation from a fuzzy enriched PAG list. In this study, we developed todenE (topology-based and density-based ensemble) clustering, pioneering in integrating topology-based and density-based clustering methods to detect PAG communities leveraging the PAG network and Large Language Models (LLM). In computational genomics annotation, the genes can be grouped/clustered through the gene relationships and gene functions via guilt by association. Similarly, PAGs can be grouped into higher-level clusters, forming concise functional representations called Super-PAGs. TodenE captures PAG-PAG similarity and encapsulates functional information through LLM, in characterizing network-based functional Super-PAGs. In synthetic data, we introduced a metric called the Disparity Index (DI), measuring the connectivity of gene neighbors to gauge clusterability. We compared multiple clustering algorithms to identify the best method for generating performance-driven clusters. In non-simulated data (Gene Ontology), by leveraging transfer learning and LLM, we formed a language-based similarity embedding. TodenE utilizes this embedding together with the topology-based embedding to generate putative Super-PAGs with superior performance in semantic and gene member inclusiveness.

## Introduction

Systems biology, when coupled with high-throughput technologies like proteomics, allows for the discovery of novel pathway biomarkers, which are essential for understanding complex diseases [1]. Functional genomics analysis and system biology are critical and fundamental for understanding the genetic basis of human health [2, 3]. It enhances our knowledge of pathogenic pathways by examining the interactions between different types of molecules, ultimately creating a comprehensive map from genotypes to phenotypes. With a deeper understanding of the molecular processes driven by complex molecular interactions and chemical reactions, biomedical discovery is expedited, leading to numerous applications in the studies of complex diseases. These include molecular risk stratification [4, 5, 6], pre-clinical drug screening [7, 8, 9], prognosis prediction [10, 11, 12], new biomarker discovery [13, 14, 15], and precision medicine [16, 17, 18]. With the advent of new and advanced techniques for multi-omics high-throughput sequencing, a vast amount of genomics data has been generated, presenting both challenges and opportunities. This data illuminates the complex interactions among genes, gene products, and the environment, ushering in a new era of molecular understanding and leading to novel hypotheses regarding the prevention and treatment of complex diseases. Integrative systems biology approaches, combined with data mining techniques, have been crucial for uncovering the role of genetic variations in diseases and mining disease-specific molecular association profiles across complex datasets [19, 20, 21, 22]. Network-based systems biology approaches have been instrumental in constructing pathway-connected networks, analyzing dose-response relationships, and prioritizing drug targets for repositioning in various diseases, from infectious diseases to cancer [23, 24, 25].

Computational functional genomics leverages computational approaches to address challenges in functional genomics, enhancing data interpretability and facilitating novel knowledge discovery. For a deeper understanding of molecular pathways and genetic control mechanisms [26], various tools and databases have been developed to perform gene set enrichment analysis by processing gene list to extract a statistical measure of shared biological features. These include Gorilla [27], DAVID [28], WebGestalt [29], EnrichR [30], PANTHER Gene List Analysis [31]), TermMapper [32], and g:profiler [33], which frequently used to provide valuable interpretation into high-throughput data. With increasing demands from multi-omics studies, several challenges arise. These include adequately covering the extensive content of multi-omics data, rendering complex network-based models, and providing advanced features for in-depth insights in integrative analysis [34]. Therefore, the integrative genesets, networks, pathways analysis (GNPA) has been introduced and leads the new height of information integration in the recently developed GNPA application, such as PyGNA [35], NDEx IQuery [36], PAGER web APP [34], and PANGEA [37]. Meanwhile, the incline of heterogeneous resources has significantly enhanced the analytical power, enabling comprehensive analysis from various sources, including pathways (Reactome [38], KEGG [39], BioCarta, WikiPathway [40], SPIKE, NCI Nature Curated pathway [41], GeoMx Cancer Transcriptome Atlas [42]), ontology annotations (gene ontology annotation and Human Phenotype Ontology [43]), tissue-specific expressions (GTEx [44], NGS Catalog [45], TCGA [46], PathologyAtlas [47], and FANTOM5 [48]), cell-specific expressions (CellAtlas [49]), disease phenotypes (GAD, GWAS Catalog [50], Phewas [51], and Online Mendelian Inheritance in Man), drug-targets (PharmGKB [52], and DSigDB [53]), and miRNA-gene interactions (Microcosm Targets [54], TargetScan [55] and mirTARbase [56]). Pathways, annotated gene lists and gene signatures (PAGs) have been introduced to standardize gene sets from heterogeneous data sources. However, the integration of multiple heterogeneous terms remains incomplete, leading to copies of gene sets in the gene set enrichment analysis and network analysis in practice [57, 58].

To address the heterogeneities and inconsistencies of Pathway Annotated Genes (PAGs), we introduce super-PAGs to represent super-clusters of PAGs and develop a new algorithm called Topology-Based and Density-Based Ensembled Clustering (Toden-E) to generate super-PAGs from regular gene sets. Toden-E features three advanced capabilities. First, Toden-E introduces a new metric, the Disparity Index for quantifying “clusterability”, indicating whether the network can be clustered into super-PAGs based on the distribution of the clustering coefficient [59]. Second, Toden-E achieves optimal performance in super-PAG mining by integrating topology-based features from PAG-PAG networks and density-based features from PAG descriptions using an ensemble model. Third, Toden-E provides summaries of the super-PAGs by synthesizing descriptions from super-PAG members using a large language model. We tested Toden-E on the gene ontology network and demonstrated that Toden-E outperforms baseline models that utilize only individual type of information in generating super-PAGs. We anticipate that super-PAGs will play a pivotal role in biology and biomedicine by collecting, standardizing, and organizing gene set knowledge to facilitate multiscale biomedical data integration and analysis.

## Methods

### Overview design of Toden-E workflow

Toden-E is developed to enhance functional genomics analysis by creating abstract biological concepts that integrate bio-semantic and biological network information. It employs a complex workflow that leverages graph transformations and machine learning models (**Fig. 1**).

**Fig. 1.**
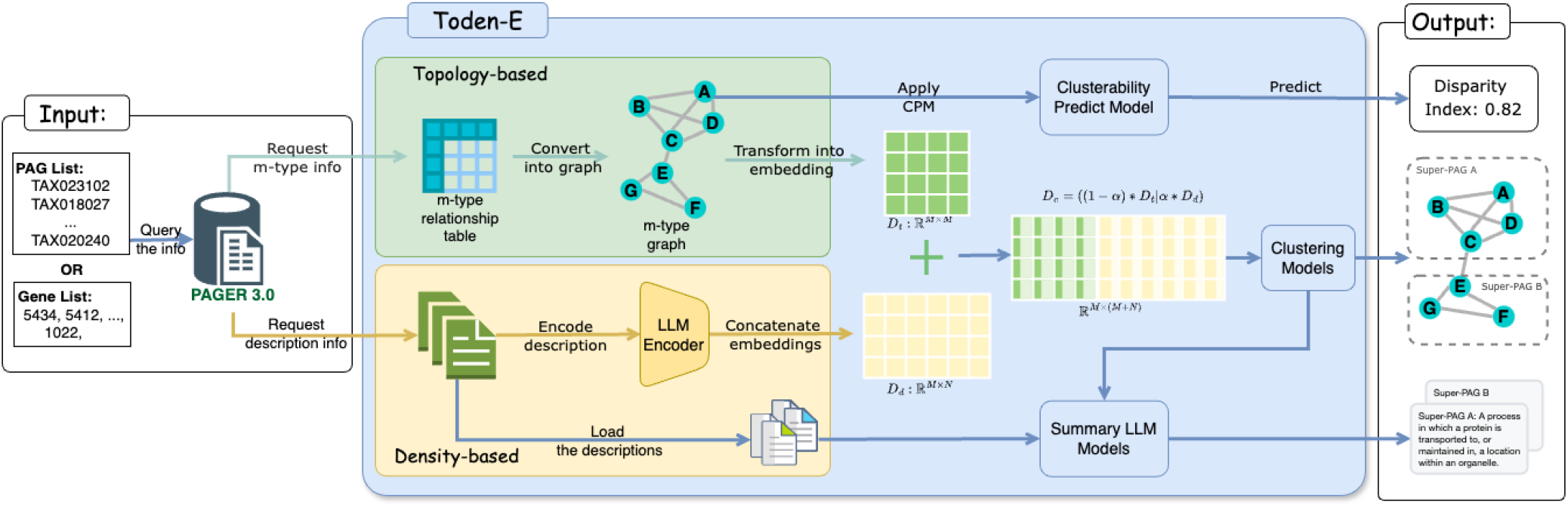
The framework of Toden-E. **Input:** The input can be either a PAG list identified by PAG IDs or a gene list identified by gene Entrez IDs. The system queries the PAGER 3.0 database for m-type PAG-PAG relationship info and PAG description info. **Topology-based:** Converts m-type relationship data into a graph, applies clustering methods to propose super-PAGs, and uses a clusterability prediction model to assess feasibility. **Density-based:** Encodes descriptions with an LLM encoder to create embeddings, which are then clustered. **Output:** Produces a Disparity Index score, super-PAG proposals, and summary descriptions for each super-PAG.

The input for Toden-E can be either PAGs or individual genes. PAGER 3.0 will retrieve co-membership-based (m-type) PAG–PAG relationships [60] and PAG descriptions for Toden-E. If a PAG list identified by PAG IDs is provided, PAGER 3.0 will directly retrieve the m-type PAG-PAG relationships and PAG descriptions. If genes are provided as input, PAGER 3.0 will perform gene set enrichment analysis to generate the significantly enriched PAGs, along with their m-type PAG-PAG relationships and descriptions.

Subsequently, Toden-E performs an ensemble clustering procedure along two parallel paths: Topology-based information processing and Density-based information processing. The details of these processes are described in the following sections. The outputs of these two processes are two embedding matrices, *D*_*t*_ and *D*_*d*_, representing the topology information and bio-semantic information, respectively. We ensemble these two types of information by concatenating the matrices and applying a hyperparameter *α* to adjust the weight of each data source, resulting in a new embedding matrix *D*_*e*_. Clustering methods are then applied to *D*_*e*_ to obtain the final clustering results, which form the putative super-PAGs. These clustering results are also used as input for summarization models to generate descriptions for each cluster as an additional output.

To improve the robustness and accuracy of Toden-E’s clustering, we integrated a consensus clustering algorithm into its workflow. Consensus clustering is well-suited for aggregating the results of multiple clustering runs to provide a stable assessment of the clustering solution, mitigating sensitivity to initial conditions. Applied to the ensemble embedding matrix *D*_*e*_, which combines topology-based and density-based information, consensus clustering helps determine the optimal number of clusters.

Toden-E generates three types of outputs: the Disparity Index to gauge clusterability, the putative super-PAGs, and a summary description for each super-PAG.

### Topology-Based Clustering Process

The m-type relationship information, which consists of PAG-PAG pairs with membership strength scores [60], was used to construct an m-type PAG-PAG network. To discover super-PAGs, we first construct a graph using the m-type relationship information. The m-type relationships quantify the significance of shared genes between pairs of PAGs using a hypergeometric distribution. To form the topology-based clusters, we applied three clustering algorithms to the m-type PAG-PAG network. The Girvan-Newman algorithm identifies communities through edge removal, the Louvain algorithm optimizes modularity, and spectral clustering utilizes the Laplacian matrix for dimensionality reduction (see Table. 1). These algorithms are instrumental in discovering super-PAGs by effectively clustering the gene set graph.

**Table 1.** Comparison of Clustering Methods.

| Method | Parameters | Scalability | Use Case | Geometry |
| --- | --- | --- | --- | --- |
| K-Means | Number of clusters | Very large $n$, medium clusters | General-purpose | Euclidean distances |
| Spectral Clustering | Number of clusters | Medium $n$, small clusters | Few clusters | Graph distance |
| Agglomerative Clustering | Clusters/threshold, linkage, distance | Large $n$ and clusters | Many clusters, non-Euclidean | Any pairwise distance |
| HDBSCAN | Minimum membership, point neighbors | Large $n$, medium clusters | Uneven sizes, outlier removal | Nearest points distances |
| Louvain | Resolution parameter | Very large $n$ | Community Detection (CD) in large networks | Graph-based modularity |
| Girvan-Newman | None (hierarchical edge betweenness) | Medium $n$ | CD in small to medium networks | Graph-based modularity |

### Converting PAG Descriptions to Latent Embedding Using SBERT

The PAG descriptions are converted into an embedding matrix for further clustering using a Large Language Model (LLM) encoder. In this study, we utilize Sentence Transformers (SBERT) [61] as the encoder.

SBERT is a modification of the pretrained BERT network designed to derive semantically meaningful sentence embeddings. Unlike BERT, which is typically used for token-level tasks, SBERT employs Siamese and triplet network structures to generate fixed-size sentence embeddings that can be compared using cosine similarity.

The embeddings generated by SBERT are then used for clustering. Various clustering algorithms, such as K-means, Agglomerative Clustering, and HDBSCAN, are applied to these embedding vectors to identify clusters of PAGs, referred to as super-PAGs. This process leverages the semantic similarities captured by SBERT to group gene sets with similar descriptions, facilitating deeper insights into the functional relationships among PAGs.

### Density-Based Clustering Process

We use the outputs of the SBERT encoder as embeddings for the descriptions of gene sets. These embeddings are then subjected to various clustering algorithms (see **Table. 1**), including K-means, Agglomerative Clustering, and HDBSCAN, to effectively cluster the data.

All the clustering algorithms utilize the SBERT latent embeddings of PAG descriptions as input feature vectors. PAGs with high description similarity are grouped together to form putative super-PAGs. Each clustering algorithm has distinct advantages. For instance, K-means excels in speed and scalability, making it suitable for handling a large number of PAGs. Agglomerative clustering offers flexibility in distance metrics and does not require specifying the number of clusters in advance. HDBSCAN provides automatic determination of clusters and effectively handles incoherent PAGs. The choice of algorithm can influence the granularity and structure of the resulting super-PAGs.

### Ensemble Method for Combining Embeddings

The ensemble method in Toden-E combines the embeddings derived from both m-type PAG-PAG network (topology-based) and description embedding (density-based) to form a unified embedding matrix, ensuring a holistic approach to clustering and summarizing gene sets in functional genomics.

To achieve this, we first compute the topology-based embedding matrix *D*_*t*_ from the m-type relationships and the bio-semantic embedding matrix *D*_*d*_ from the SBERT-encoded descriptions. These matrices are then combined to form a new embedding matrix *D*_*e*_:

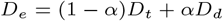

where *α* is a hyperparameter that adjusts the weight of the two data sources, allowing for scalable integration based on the relative importance of structural versus content-based information.

The combined embedding matrix *D*_*e*_ is then subjected to clustering algorithms, such as K-means, Agglomerative Clustering, or HDBSCAN, to discover super-PAGs. These clustering results are also utilized for summarization, providing concise and informative descriptions for each super-PAG.

In the advanced mode, the embedding matrix *D*_*e*_ can be extended with additional features of gene sets, such as expression levels, mutation data, or other relevant biological information, to further enhance the discovery of super-PAGs.

### Evaluation of Clustering Algorithms Using ARI

The Adjusted Rand Index (ARI) is a critical metric for evaluating the similarity between two clustering results, adjusting for chance. Unlike the Rand Index, which does not account for random agreement, ARI provides a more accurate measure of clustering quality.

### Synthetic Network Modeling

To explore the inherent clusterability of graphs, we synthesize networks utilizing the stochastic block model (SBM) by varying the intra-cluster and inter-cluster connection probabilities. We exhaustively generate combinations of intra-cluster and inter-cluster connection probabilities in increments of 0.1, ranging from 0.1 to 1.0, for two distinct clusters. These synthetic network models will be used for training the clusterability predictive model and generate the Disparity Index.

### Clusterability Assessment

To assess the conceptual clusterability of a given network, Toden-E incorporates a predictive model trained on synthetic network datasets. Since clusterability is challenging to measure directly, Toden-E uses synthetic networks to generate cluster coefficient distributions and learn to predict the validity of clustering algorithms using Adjusted Rand Index (ARI). We employ four commonly used classification models, each evaluated using various metrics, including accuracy, precision, recall, and F1-score, to identify the best model for predicting clusterability.

#### Algorithm 1

An algorithm to develop a prediction model to determine the clusterability of a graph.

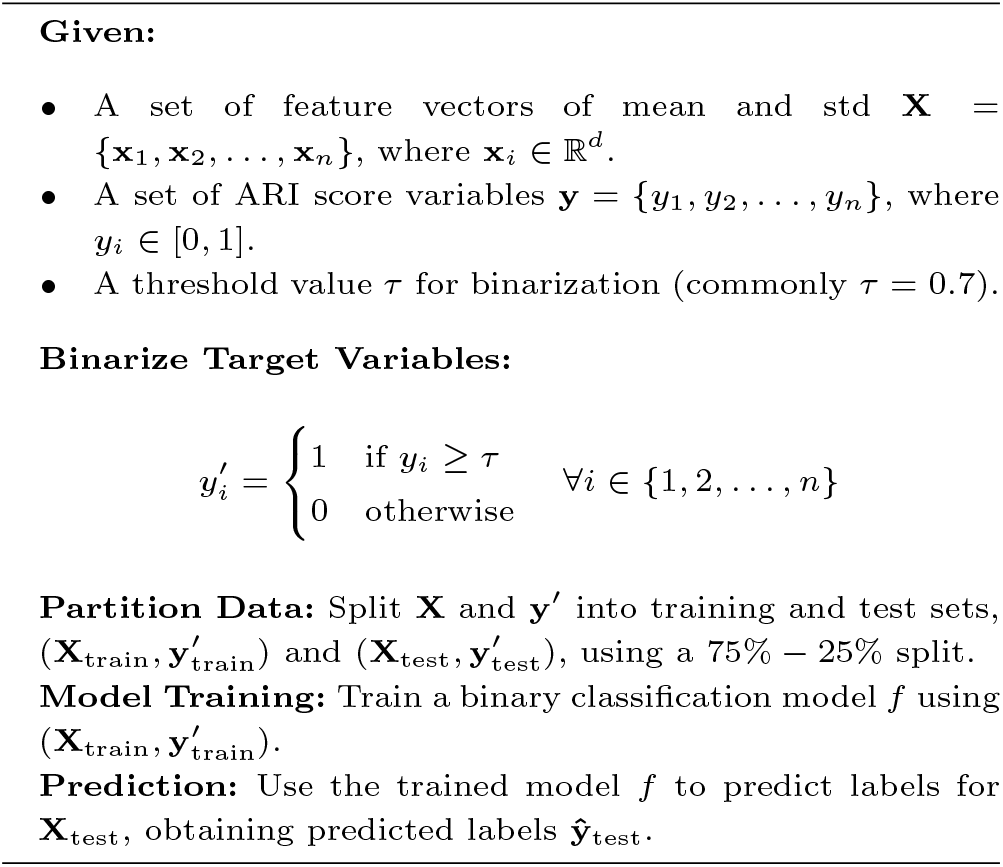

### Gene Ontology Annotation Network in PAGER 3.0 as Benchmarking Dataset

To assess the practical applicability of Toden-E, we employed the Gene Ontology Annotation (GOA) m-type PAG-PAG network derived from the GOA database [62]. In this network, the ground truth relationships are represented by the “is-a” hierarchical relationships, which define the parent-child connections between GO terms. Toden-E utilizes the m-type GOA-GOA relationships and their accompanying descriptions as input to infer and recover these “is-a” relationships.

For this study, we extracted an “is-a” relational network from GO terms classified under “biological processes.” To ensure a rigorous and fair evaluation, we applied two key selection criteria: (1) the GO terms must fall under distinct parent terms to capture diverse biological concepts, (2) the GO terms must not share a common ancestor, thereby maintaining conceptual independence, and (3) the number of nodes in a given PAG should be greater than 10 and less than or equal to 100. These groups are termed SPAGs 10-100. Based on these criteria, two datasets were constructed: the first containing 127 records, each consisting of a pair of GO terms, and the second comprising 300 records, each involving a triplet of GO terms.

After analyzing the distribution of nodes that met the first two criteria, we also developed an additional benchmark where the number of nodes is greater than 5 but less than or equal to 10 which is termed SPAGs 5-10.

### Utilizing Large Language Models for PAG Summarization

In our study, we harnessed the power of Large Language Models (LLMs) to enhance the interpretability of Pathways, Annotated Gene lists, and Gene Signatures (PAGs) by summarizing their descriptions. Given the often extensive and detailed functional descriptions associated with each PAG, manually synthesizing these into concise summaries for higher-level clusters, or Super-PAGs, can be laborious and inefficient. To address this, we employed LLMs to automate the summarization process, ensuring that each Super-PAG is represented by a clear, informative sentence.

We first concatenated all functional descriptions associated with each PAG within a cluster into a single, comprehensive text block, ensuring that the language model (LLM) receives full contextual information about the cluster’s biological functions. Using this concatenated text, the LLM generated a summary by processing the input and distilling it into a coherent sentence that encapsulates the essence of the cluster’s biological functions. This summary provides a high-level overview of the Super-PAG, facilitating easier interpretation by researchers. Given the inherent input size limitations of LLMs, we optimized our method by monitoring the length of the concatenated descriptions. When the input approached the model’s capacity, we applied additional summarization steps to further condense the text before feeding it into the LLM. This process ensured that all relevant information was retained while staying within the model’s input limits.

## Results

### Topology-Based Clustering Performance of Synthetic Network

To evaluate the effectiveness of topology-based clustering methods, we constructed a Stochastic Block Model (SBM) network with various settings for the probabilities of inter-cluster and intra-cluster connections. Initially, we set the probability of inter-cluster connections at 0.5 and intra-cluster connections at 0.1 to test the performance of three main clustering algorithms: the Girvan-Newman Algorithm, the Louvain Algorithm, and Spectral Clustering.

We further extended our analysis by applying these algorithms to SBM networks with different probability settings: (inter-cluster vs. intra-cluster) as (0.3, 0.1), (0.5, 0.01), (0.6, 0.4), and (0.8, 0.1) by observing the results across these varying configurations (see **Fig. S1**), we found that Spectral Clustering consistently outperformed the other two algorithms in accurately detecting the underlying cluster structures.

Given its superior performance across different network settings, we selected Spectral Clustering as the primary method for analyzing synthetic datasets in the subsequent sections of our study. This choice ensures robust and reliable identification of community structures, especially in networks characterized by complex modularity and connectivity patterns.

### Exploring Clusterability: The Performance of Clusterability Predictive Model

The clusterability predictive model was designed to classify synthetic networks into unclusterable and clusterable categories with an accuracy of 0.88 and an F1 score of 0.9.

The ARI was utilized as the standard for gauging clusterability in the synthetic networks. We employed spectral clustering, the best-performing algorithm reported in the topology-based clustering, to predict clusters and evaluate their agreement with the ground truth clusters. As indicated in the ARI measurement heatmap shown in **Fig. 2**, a distinct boundary emerged, demarcating networks that could be correctly clustered from those that could not. We set the ARI threshold at 0.7 (**Fig. 3a, 3b, and 3c**), transforming the network clusterability assessed by spectral clustering into a binary classification problem.

**Fig. 2.**
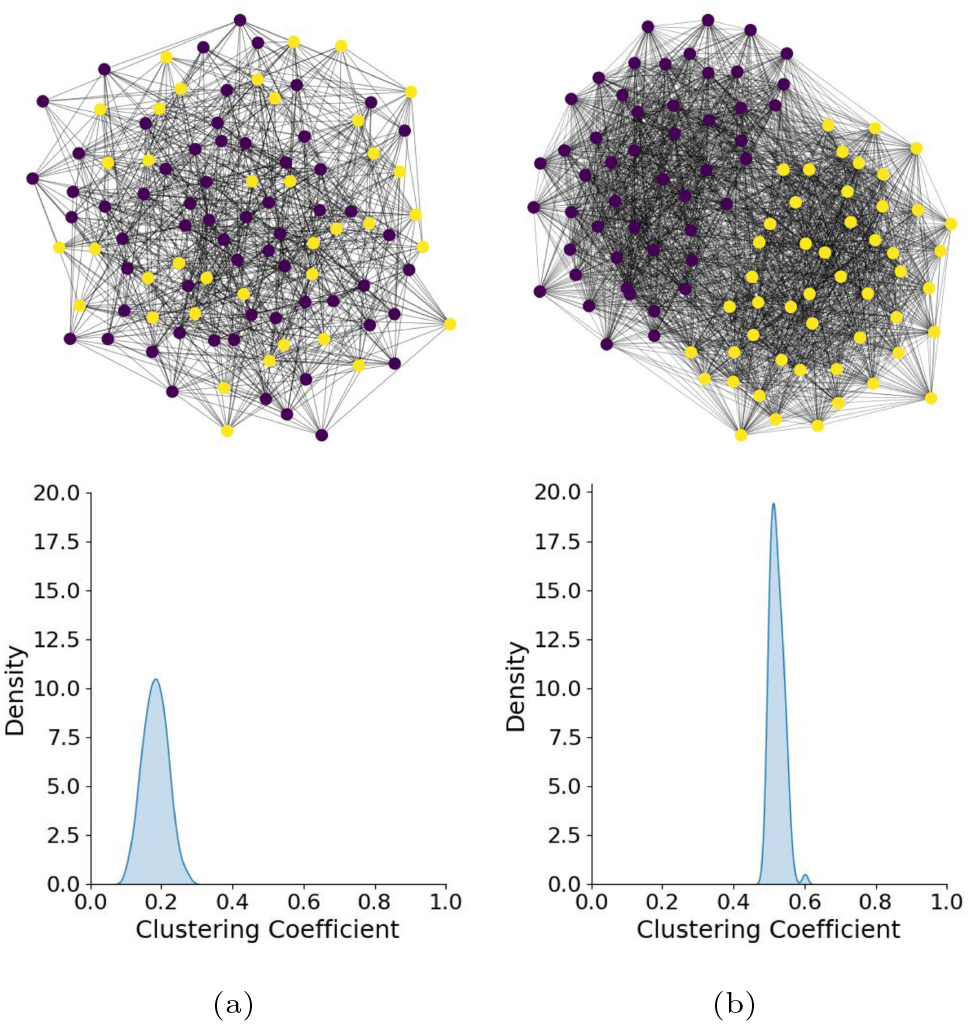
Comparison of the clustering coefficient distribution between two synthetic networks: one clusterable and the other non-clusterable.(a) Network graph generated by the stochastic block model (SBM) with an intra-cluster connection probability of 0.2 and an inter-cluster connection probability of 0.2. Its clusterability is low. (b) Network graph generated by the stochastic block model (SBM) with an intra-cluster connection probability of 0.7 and an inter-cluster connection probability of 0.3. Its clusterability is high.

**Fig. 3.**
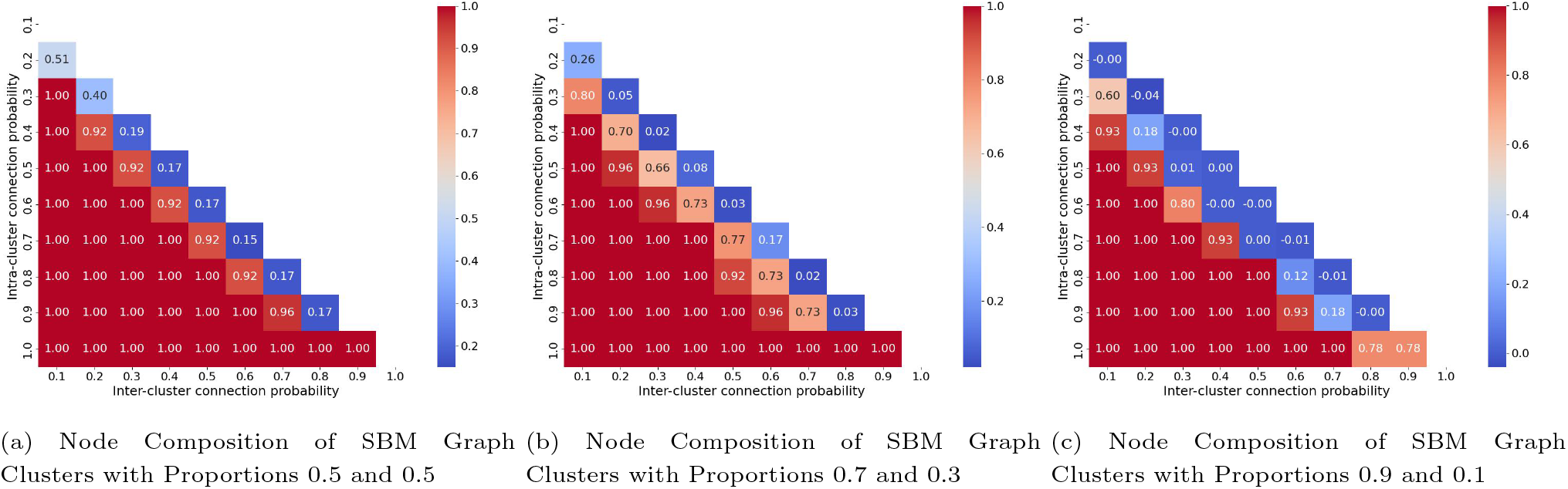
Heatmap of Adjusted Rand Index (ARI) Comparing Clusterability Across Three Settings. It illustrates the distinct clusterability variations when using spectral clustering on graphs generated by a stochastic block model (SBM) with three different configurations. Each configuration’s ARI score is visualized as a heatmap, clearly delineating the differences in clustering performance.

We developed a clusterability predictive model described in **Algorithm 1**. The input of the model is the mean and variance of cluster coefficient distribution and the output is the binary outcome network clusterability. In **Fig. 2**, we can intuitively identify the differences in cluster coefficient distribution between unclusterable and clusterable networks. The unclusterable network displayed a normal distribution with low cluster coefficients (0.2). In contrast, the clusterable network exhibited a high cluster coefficient (0.5 to 0.6) with a bimodal distribution. Particularly, we employed Logistic Regression, Support Vector Machines (SVM), Random Forest, and K-Nearest Neighbors (KNN) models. The Random Forest model demonstrated superior performance with an F1-score of 0.90, signifying a high balance between precision and recall (**Table. 2**). Logistic Regression also showed commendable results, particularly in terms of recall.

**Table 2.** Comparison of different methods based on performance metrics.

| Methods | Accuracy | Precision | Recall | F1 |
| --- | --- | --- | --- | --- |
| Logistic Regression | 0.76 | 0.73 | 1.00 | 0.84 |
| SVM | 0.67 | 0.66 | 1.00 | 0.79 |
| Random Forest | **0.88** | 1.00 | 0.81 | **0.90** |
| KNN | 0.76 | 1.00 | 0.63 | 0.77 |

Overall, the clusterability predictive model offers a quantitative method for evaluating the validity of clustering algorithms in network clusterability problems. This approach is especially valuable for assessing complex network structures where discerning clear clusterability boundaries is critical.

### Topology-Based Clustering Performance Analysis

In our analysis of topology-based clustering, we evaluated the effectiveness of various clustering methods in identifying structural communities within synthetic datasets. The methods tested included Louvain, Spectral Clustering, and Agglomerative Clustering. The Adjusted Rand Index (ARI) was used as the primary metric to assess clustering performance across two datasets: a Two-Groups Dataset and a Three-Groups Dataset.

As depicted in **Fig. 4**, Spectral Clustering consistently outperformed the other algorithms, particularly in the Three-Groups Dataset, where it achieved the highest ARI scores. Louvain Clustering also demonstrated strong performance, especially in the Two-Groups Dataset, but showed more variability in the Three-Groups Dataset. On the other hand,

**Fig. 4.**
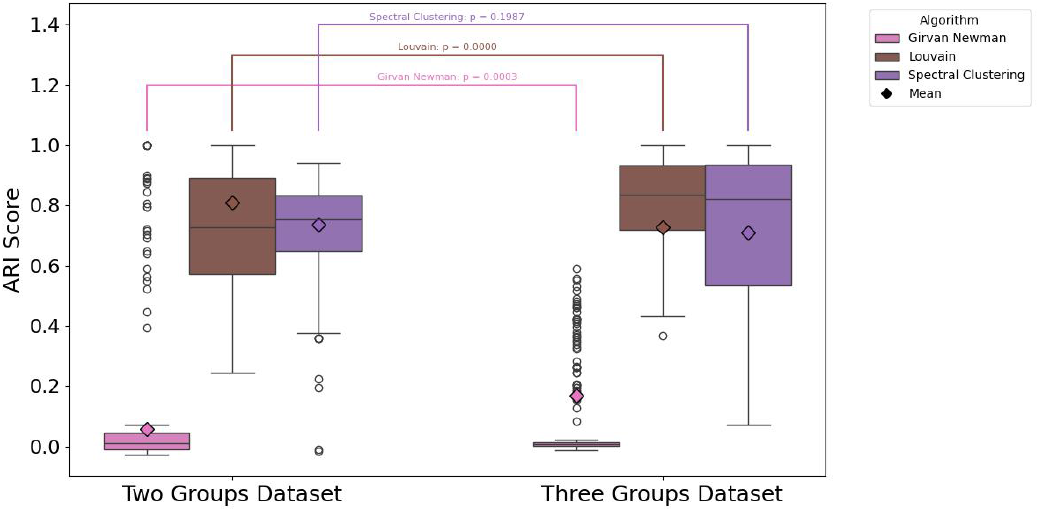
Performance of the Topology-based model on the evaluation dataset SPAGs 10-100.

Agglomerative Clustering exhibited the most variability, with notably lower ARI scores in the Three-Groups Dataset, indicating that it may be less effective in handling more complex cluster structures. The statistical significance of these differences was confirmed by p-values, highlighting the superior performance of Spectral Clustering across different dataset complexities.

In addition to analyzing the SPAGs 10-100 group, we also applied the same clustering analysis to SPAGs 5-10. Interestingly, the performance was even better for SPAGs 5-10, as clustering a lower number of nodes is generally more straightforward compared to clustering a larger number of nodes. Given the inherent simplicity of smaller clusters, the clustering algorithms performed more consistently with higher accuracy in this group. Nevertheless, we present the performance on SPAGs 10-100 as our main results, since these larger clusters provide a more challenging and informative benchmark for evaluating the robustness of the clustering methods.

Based on these results, Spectral Clustering was chosen as the preferred method for further analysis due to its consistent and reliable performance in accurately identifying topology-based clusters, even in more complex scenarios.

### Density-Based Clustering Performance Analysis

**Fig. 5** illustrates the performance comparison of K-Means, Agglomerative Clustering, and HDBSCAN algorithms on two synthetic datasets: a Two-Groups Dataset and a Three-Groups Dataset, using the Adjusted Rand Index (ARI) as the evaluation metric. In the Two-Groups Dataset, both K-Means and Agglomerative Clustering performed similarly, achieving high ARI scores, while HDBSCAN showed greater variability with a broader distribution of ARI scores, indicating less consistent clustering performance. In the Three-Groups Dataset, Agglomerative Clustering maintained high ARI scores, suggesting its robustness in more complex clustering tasks. However, HDBSCAN’s performance significantly dropped, as evidenced by the lower median ARI score and wider spread, reflecting its challenges in accurately clustering more complex datasets. K-Means, while slightly less effective than Agglomerative Clustering, still demonstrated relatively stable performance across both datasets. These results suggest that Agglomerative Clustering is the most reliable method among the three, particularly in scenarios involving multiple, well-separated clusters.

**Fig. 5.**
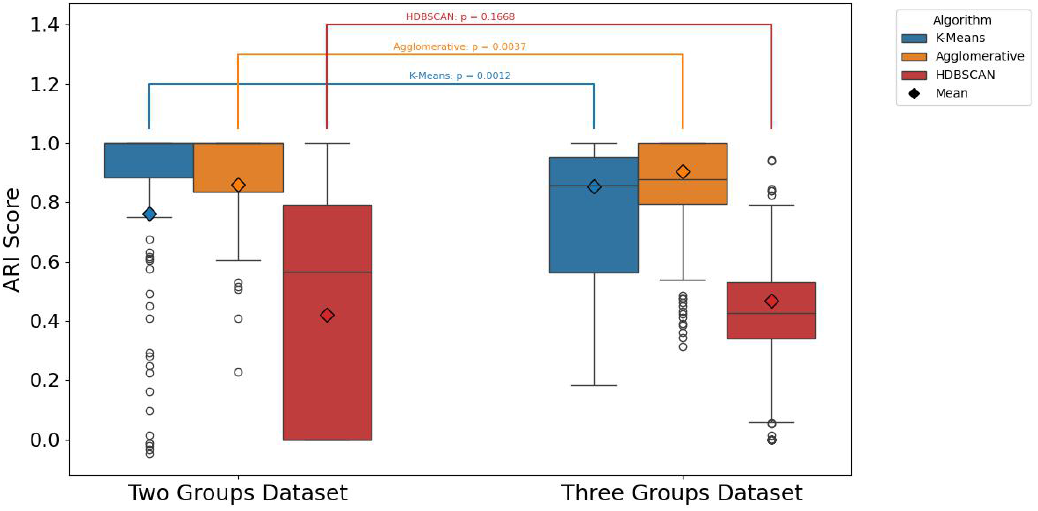
Performance of the Density-based model on the evaluation dataset SPAGs 10-100.

### Toden-E Outperforms Individual Method

Toden-E demonstrated superior performance in recovering “is-a” relationships in GOA networks compared to individual topology-based and density-based clustering methods. After applying the three clustering algorithms to the GOA benchmarking datasets, the Toden-E model with a weighted ensemble approach showed higher ARI scores. We tested Toden-E in both regular cases, with the average GOA size being 5 (the average size of GOA), and in extreme cases, where the GOA size varied from 10 to 100. The results indicated that Toden-E effectively managed different GOA sizes, showcasing its robustness and adaptability in various scenarios. In the regular GOA dataset, the ARI achieved its highest score with minimal variance among the three algorithms when the coefficient *α* was set to 0.6. This indicates that a 40% weighted topology-based embedding concatenated with a 60% weighted density-based embedding resulted in optimal clustering performance (see **Fig. 6**).

**Fig. 6.**
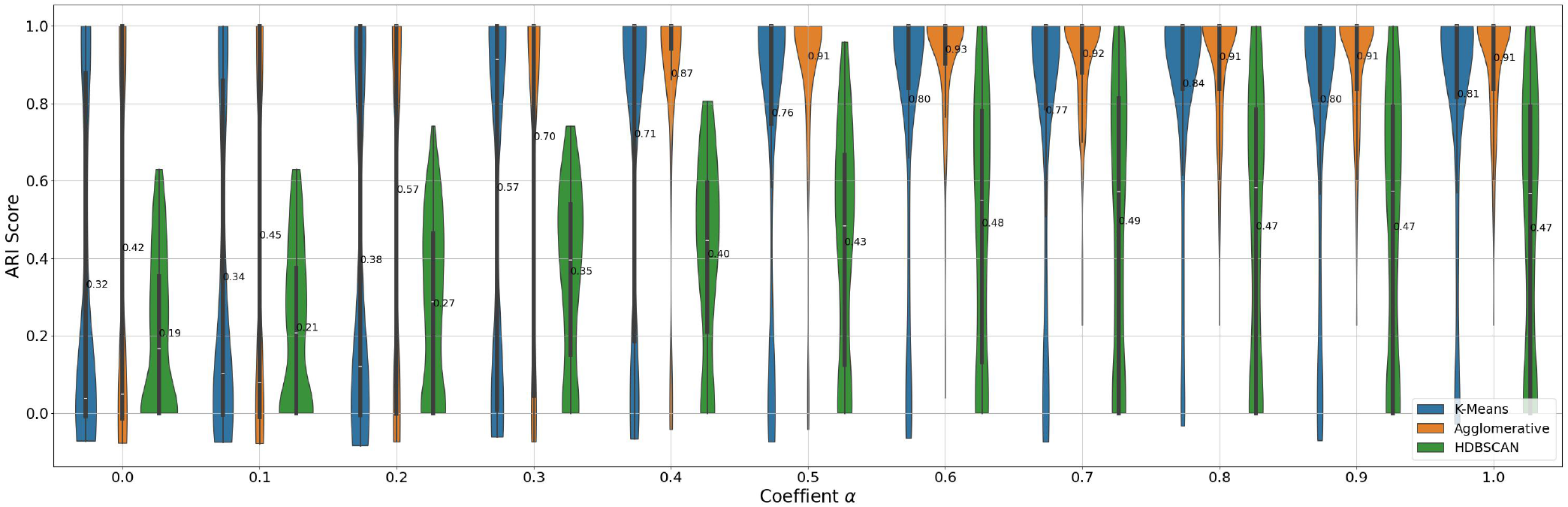
Performance of the TodenE model across various coefficient *α* settings, combining topology-based and density-based methods.

### The PAG Summarization Captures Important Information in the Benchmarking Dataset

In this study, we employed two models—BART and GPT-2—for summarizing PAG descriptions. BART, a model specifically designed for summarization tasks, proved to be highly effective in generating concise and coherent summaries, while GPT-2, a generative text model, required more careful prompting to summarize concatenated descriptions. For the SPAGs 5-10 dataset, BART achieved a mean similarity score of 0.63, demonstrating its capability in capturing key information from biological process descriptions. This result underscores the potential of language models like BART for simplifying complex biological data into digestible formats.

Looking ahead, we believe that the summarization performance can be further improved by integrating more advanced language models, such as Chat-GPT 4 or LLaMA 3, which are better suited to capture subtle semantic details. However, it is important to note that utilizing these advanced models would require addressing their significant computational demands.

### Case Study of Chronic Myeloid Leukemia Tyrosine Kinase Inhibitor Resistant vs. Sensitive

Toden-E was exemplified through a case study involving GOA PAGs derived from gene set enrichment analysis of differentially expressed genes in Chronic Myeloid Leukemia (CML) samples resistant and sensitive to Tyrosine Kinase Inhibitors (TKIs) [63][Toden-E explored two major GOA clusters within these PAGs. One cluster summarized the concepts related to the development of resistance to chemotherapy drugs, rendering the treatment ineffective in inhibiting cancer cell proliferation or inducing cell death. Conversely, the other cluster consisted of sensitive cells that continued to respond favorably to therapeutic agents. Additionally, by retrieving the “is-a” relationships to construct the GOA tree, the two major GOA clusters were recovered in different branches of the tree (**Fig. 7**). This real-world application of our framework highlights its potential utility in distinguishing differential cellular responses to treatment in leukemia.

**Fig. 7.**
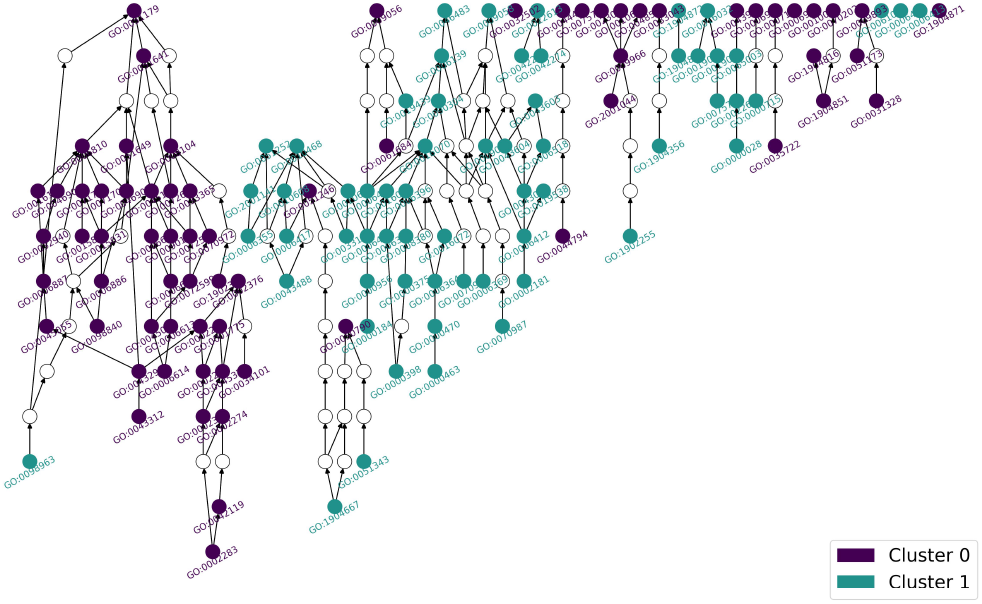
To visualize the clusters formed by TodenE from the Leukemia gene sets, 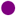 (purple circles) and 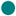 (green circles) represent two distinct clusters. White circles serve a specific function in this representation: they are placeholders used to maintain the integrity of the tree structure, which originates from the “is-a” relationships found within the GOA dataset [62].

## Conclusion and Discussion

We demonstrated Toden-E as a novel way of capturing and representing biologically and functionally important super-PAGs from regular PAGs. With multi-omics data being routinely generated, the results are all contributing to the increased diversity of PAGs in the PAGER database. It is critical to perform Toden-E to ensure that super-PAGs clearly and comprehensively represent the results of GNPA. We introduced a critical metric, the clustering coefficient distribution index, for gauging “Clusterability” in PAG networks and evaluated the super-PAG construction using the stochastic block model. The cluster coefficient metric enables the evaluation of any type of network to be clustered or not and further provides guidance for further refining networks with a higher edge confidence cutoff. As the complexity of the PAG content integration increases while considering the PAG-PAG network and PAG semantic description, Toden-E implements the ensembled embeddings from the m-type PAG-PAG relationship and the PAG description to balance the knowledge-based information and gene memberships to achieve the best performance. Toden-E will be flexible in integrating additional information layers, such as embeddings from gene-miRNA relationships, gene set functional inference similarity, etc. To enhance the interpretability of super-PAGs, we performed description extraction and summarization of gene sets with state-of-the-art LLM techniques.

By combining multiple sources of information, Toden-E offers a powerful and flexible framework for functional genomics analysis, facilitating deeper insights and more accurate interpretations of gene set relationships. Particularly, the density-based clustering outcomes were analyzed to assess the effectiveness of the LLM-based feature representation. The results revealed that the embeddings generated by the SBERT model enhanced clustering quality, as indicated by improved cluster coherence and separation. Thus, Toden-E provided deeper insights into the relationships between gene sets and offered a more nuanced understanding compared to traditional clustering methods based on simple numerical features. The integration of linguistic features through LLMs, therefore, holds significant promise for advancing the interpretability and accuracy of bioinformatics analyses.

Toden-E can be extensively used for any gene set summarization based on multi-omics data and single-cell data. The representation of the gene set can be enriched by considering different types of activities from various omics sources, such as mutation burden [65], transcriptional activity [66], proteomics activity [67], and metabolic activity [68], facilitating deeper insights and more comprehensive interpretations of gene sets and putative gene set clusters. All gene set activities can be converted into embeddings to form different super-PAGs for different levels of integration. Additionally, with the prevalence of single-cell data analysis, cell-type-specific and cluster-specific super-PAGs can be generated using cell-based multi-omics profiles [69]. Therefore, Toden-E has the potential for broad data implementation and plays a pivotal role in examining non-redundant and comprehensively condensed information in GNPA, making it extremely useful for diverse applications in multi-omics and single-cell studies.

For best practices in GNPA with super-PAGs, Toden-E is poised for potential improvements. In topology-based clustering, the m-type PAG-PAG relationships were used to generate topology-based embeddings. Using a higher threshold can result in better pattern representation but may lead to a tradeoff with a lower network coverage rate, leaving some PAGs uncovered in a PAG-PAG network. One solution could be locally adaptive smoothing with signal denoising [70]. Additionally, the complexity of the network modeling increases when integrating multi-modality relationships, such as transcriptional regulatory relationships and post-transcriptional regulatory relationships. Co-embedding of PAG-PAG relationships, PAGs, and nodes can be implemented to model multi-dimensional edge features [71]. In the density-based clustering of text embeddings from PAG descriptions using LLM, SBERT can be substituted with other LLMs, including ChatGPT [72] and LLM2Vec [73]. Currently, super-PAGs are regarded as flattened one-level structures above regular PAGs. For multi-omics integration, the topology-based and density-based embeddings can be enhanced through matrix addition when dimensions are the same. The potential benefit of this approach is that setting the coefficient to 0 or 1 can yield exact results from the single embedding. In future work, Toden-E will be implemented to recover or construct multi-level architectures using auto-weighted embedding ensemble and learning.

Overall, Toden-E takes advantage of network biology and LLM to generate super-PAGs with enhanced interpretability and quality for GNPA. It shows potential in comprehensive data integration and knowledge abstractive summarization. We expect that super-PAGs generated from Toden-E will become an extensive and publicly accessible source of non-redundant gene sets with curated biological context. Meanwhile, Toden-E will serve as a powerful tool to improve and enhance the biologists’ experience in GNPA.

## Authors Contributions

Q.L.: methodology, producing results, writing the original draft and reviewing and editing. C.N: reviewing. R.W.: data collection and curation, reviewing and editing. J.Y.C.: funding acquisition, reviewing, and editing. W.K.: supervision, reviewing, and editing. Z.Y.: funding acquisition, conceptualization, supervision, methodology, writing the original draft, reviewing, and editing. All authors contributed to the article and approved the submitted version.

## Acknowledges

We acknowledge the computational support from the staff who manage the Easley HPC at Auburn University.

## Funding Statement

ZY and JYC acknowledge the grant from the University of Alabama at Birmingham Center for Clinical and Translational Science Pilot Award (UM1TR004771).

## Conflict of Interests

The authors declare that the research was conducted in the absence of any commercial or financial relationships that could be construed as a potential conflict of interest.

## Notes

### Competing Interest Statement

The authors have declared no competing interest.

