## supplementary for "Toden-E: Topology-Based and Density-Based Ensembled Clustering for the Development of Super-PAG in Functional Genomics using PAG Network and LLM"

### Evaluation Methods

Mathematically, ARI is defined as follows:

$$\text{ARI} = \frac{\sum_{ij}\binom{n_{ij}}{2} - \left[\sum_i \binom{a_i}{2} \sum_j \binom{b_j}{2}\right] / \binom{n}{2}}{\frac{1}{2}\left[\sum_i \binom{a_i}{2} + \sum_j \binom{b_j}{2}\right] - \left[\sum_i \binom{a_i}{2} \sum_j \binom{b_j}{2}\right] / \binom{n}{2}} \quad (1)$$

where $n_{ij}$ is the number of elements in both cluster $i$ in one clustering and cluster $j$ in the other clustering, $a_i$ is the sum of the elements in cluster $i$, and $b_j$ is the sum of the elements in cluster $j$. The denominator normalizes the index by accounting for the expected similarity of all pairwise combinations.

This metric provide a robust evaluation of clustering algorithms, ensuring that the results are both accurate and statistically significant. By incorporating ARI and NRI into the Toden-E framework, we achieve a thorough assessment of clustering methods across synthetic and benchmark datasets, such as gene ontology annotations (GOA) [62], leading to more reliable and interpretable bioinformatics analyses.

### The Clustering Algorithms Used for Density-Based Method

- **K-means Clustering** K-means clustering aims to partition the dataset into $K$ clusters by minimizing the within-cluster variance. The algorithm operates as follows:

  1. Initialize $K$ cluster centroids randomly.
  2. Assign each data point $x_i$ to the nearest cluster centroid $c_j$:
  $$\text{argmin}_j \|x_i - c_j\|^2$$
  3. Update the cluster centroids based on the mean of the assigned points:
  $$c_j = \frac{1}{|C_j|}\sum_{x_i \in C_j} x_i$$
  4. Repeat steps 2 and 3 until convergence.

- **Agglomerative Clustering** Agglomerative clustering is a hierarchical clustering method that builds nested clusters by successively merging or splitting them. The algorithm operates as follows:

  1. Initialize each data point as its own cluster.
  2. Compute the distance between all pairs of clusters using a linkage criterion (e.g., single, complete, average linkage):
  $$d(C_i, C_j) = \text{linkage}(C_i, C_j)$$
  3. Merge the pair of clusters with the smallest distance.
  4. Repeat step 2 until the desired number of clusters is obtained.
  5. Common linkage criteria include:

     – Single linkage (minimum distance):
     $$d(C_i, C_j) = \min_{x \in C_i, y \in C_j} \|x - y\|$$
     – Complete linkage (maximum distance):
     $$d(C_i, C_j) = \max_{x \in C_i, y \in C_j} \|x - y\|$$
     – Average linkage:
     $$d(C_i, C_j) = \frac{1}{|C_i||C_j|}\sum_{x \in C_i}\sum_{y \in C_j} \|x - y\|$$

- **HDBSCAN (Hierarchical Density-Based Spatial Clustering of Applications with Noise)** HDBSCAN is an extension of DBSCAN that converts it into a hierarchical clustering algorithm and then extracts a flat clustering based on the stability of clusters. The algorithm operates as follows:

  1. Compute the mutual reachability distance between all pairs of points:
  $$d_{\text{mreach}}(x_i, x_j) = \max(\text{core}_k(x_i), \text{core}_k(x_j), d(x_i, x_j))$$
  where $\text{core}_k(x)$ is the core distance of $x$, defined as the distance to its $k$-th nearest neighbor.
  2. Construct the minimum spanning tree (MST) of the mutual reachability distances.
  3. Condense the MST into a hierarchy of clusters based on the minimum cluster size.
  4. Extract the flat clustering by selecting the most stable clusters from the hierarchy.

- **Girvan-Newman Algorithm** The Girvan-Newman algorithm detects communities by iteratively removing the edge with the highest betweenness centrality until the graph is divided into disconnected components. The betweenness centrality $C_B(e)$ of an edge $e$ is defined as:
$$C_B(e) = \sum_{s \neq t \neq e} \frac{\sigma(s, t|e)}{\sigma(s, t)}$$
where $\sigma(s, t)$ is the total number of shortest paths from node $s$ to node $t$, and $\sigma(s, t|e)$ is the number of those paths that pass through edge $e$.
- **Louvain Algorithm** The Louvain algorithm is an iterative method that optimizes the modularity of the partition of the graph. Modularity $Q$ is defined as:
$$Q = \frac{1}{2m}\sum_{i,j}\left[A_{ij} - \frac{k_i k_j}{2m}\right]\delta(c_i, c_j)$$
where $A_{ij}$ is the adjacency matrix of the graph, $k_i$ and $k_j$ are the degrees of nodes $i$ and $j$, $m$ is the number of edges, and $\delta(c_i, c_j)$ is 1 if nodes $i$ and $j$ are in the same community, and 0 otherwise.
- **Spectral Clustering** Spectral clustering uses the eigenvalues of the Laplacian matrix $L$ of the graph to perform dimensionality reduction before applying k-means clustering. The Laplacian matrix is defined as:
$$L = D - A$$
where $D$ is the degree matrix and $A$ is the adjacency matrix. The algorithm proceeds by computing the first $k$ eigenvectors of $L$ (corresponding to the $k$ smallest eigenvalues) and using them as the new representation of the nodes. These representations are then clustered using k-means.

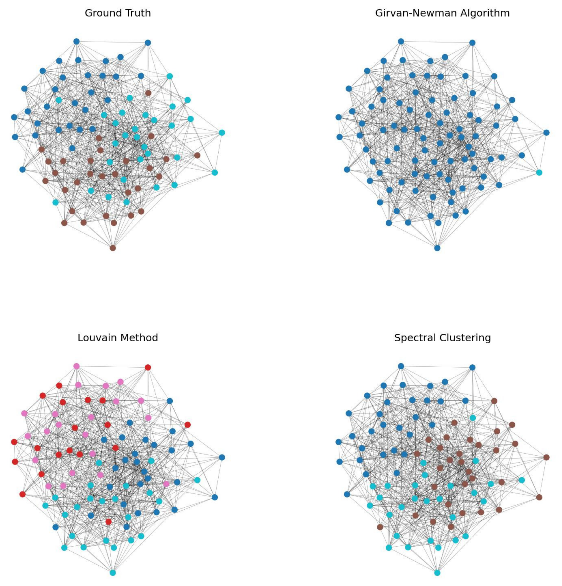


**Fig. S1.** Visualization of Ground Truth and Clustering Results in SBM Network (Intra-Probability = 0.1, Inter-Probability = 0.3). This figure compares the ground truth clusters of the synthetic SBM network with clustering results from three algorithms: Girvan-Newman, Louvain, and Spectral Clustering. Spectral Clustering shows the closest match to the ground truth, highlighting its effectiveness in accurately identifying network structures under the given probability settings.

### Topology-Based Clustering Performance on SPAGs 5-10

We applied topology-based clustering methods, including Louvain, Spectral Clustering, and Agglomerative Clustering, to the SPAGs 5-10 group. The Adjusted Rand Index (ARI) was used to evaluate the effectiveness of these algorithms. As shown in **Fig. S2** Spectral Clustering consistently outperformed the other methods, particularly in the Three-Groups Dataset, where it achieved the highest ARI scores ($p = 0.0003$). Louvain performed well in the Two-Groups Dataset but showed more variability in the Three-Groups Dataset, with non-significant differences ($p = 0.4764$). Agglomerative Clustering, although effective in simpler structures, exhibited the most variability and lower ARI scores, particularly in more complex datasets ($p = 0.005$). These results reinforce that Spectral Clustering is better suited for identifying clusters in smaller node sets, such as SPAGs 5-10, where the overall performance is enhanced compared to larger node sets.

### Density-Based Clustering Performance on SPAGs 5-10

**Figure S3** presents the performance of K-Means, Agglomerative Clustering, and HDBSCAN on SPAGs 5-10, evaluated using the Adjusted Rand Index (ARI). In the Two-Groups Dataset, K-Means and Agglomerative Clustering performed similarly, both achieving high and consistent ARI scores. HDBSCAN, on the other hand, exhibited greater variability, with a broader range

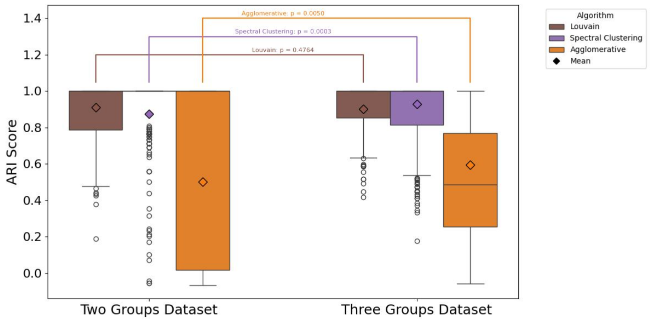


**Fig. S2.** Performance of the Topology-based model on the evaluation dataset SPAGs 5-10.

of ARI scores, reflecting its less consistent performance in this simpler scenario.

In the Three-Groups Dataset, Agglomerative Clustering continued to demonstrate strong performance, maintaining high ARI scores, and proving its robustness in more complex clustering tasks. K-Means showed relatively stable performance but was slightly less effective than Agglomerative Clustering. HDBSCAN, however, experienced a significant drop in performance, as evidenced by its lower median ARI score and a wider distribution, indicating challenges in accurately clustering datasets with more complex structures.

These results suggest that while Agglomerative Clustering is the most reliable method for SPAGs 5-10, particularly when dealing with multiple well-separated clusters, K-Means still provides relatively stable results. HDBSCAN, though useful in specific cases, may struggle with consistency in more complex clustering scenarios.

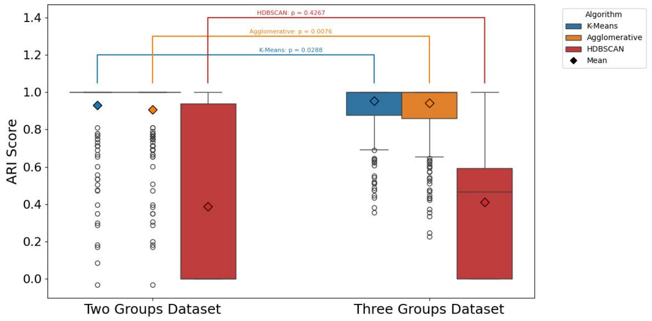


**Fig. S3.** Performance of the Density-based model on the evaluation dataset SPAGs 5-10.

### Applying Consensus Clustering to Toden-E for Optimal Cluster Number Selection

In order to enhance the robustness and accuracy of Toden-E's clustering outputs, we integrated a consensus clustering [74] algorithm into its workflow. Consensus clustering is a methodology designed for class discovery and clustering validation, particularly suited for analyzing complex biological data such as gene expression profiles. By aggregating the results of multiple runs of a clustering algorithm, consensus clustering provides a stable and reliable assessment of the clustering solution, mitigating the sensitivity to initial conditions that often affects traditional clustering methods.

For Toden-E, the consensus clustering algorithm was applied to the ensemble embedding matrix $D_e$, which combines topology-based and density-based information. By running various clustering algorithms, such as K-means and model-based Bayesian clustering, multiple times with random restarts, consensus clustering allowed us to identify the most stable and meaningful clusters. This approach not only helps in determining the optimal number of clusters but also provides insights into cluster membership stability and boundaries.

The application of consensus clustering to Toden-E ensures that the final putative super-PAGs are biologically meaningful and statistically robust. It also facilitates the visualization and interpretation of clustering results, providing researchers with a powerful tool to explore abstract biological concepts that integrate both bio-semantic and biological network information. By leveraging consensus clustering, Toden-E is better equipped to propose the best cluster number for a given set of genes or PAGs, thereby enhancing the overall utility and reliability of the method in functional genomics analysis. (see **Fig. S4**)

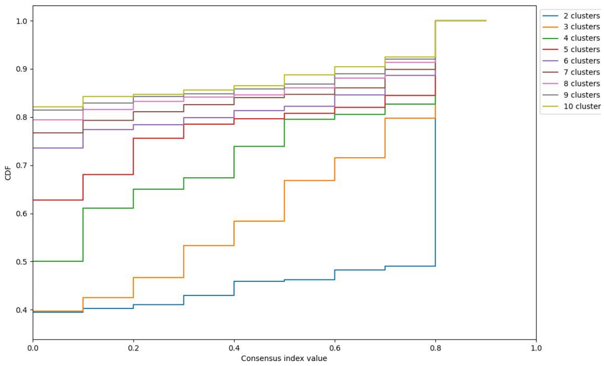


**Fig. S4.** CDF Plots from Consensus Clustering to Determine Optimal Cluster Number. It presents the Cumulative Distribution Function (CDF) plots corresponding to consensus matrices generated for different numbers of clusters (K = 2, 3,..., 10). The analysis indicates that the best number of clusters is 2, as suggested by the CDF plot with the most stable and highest consensus index.

### The Process of Converting the Description into the Embeddings

The process works as follows:

1. **Input Sentences**: Given a description $d_i$ for a gene set $G_i$, SBERT processes the text to generate an embedding vector.
2. **Encoding**: SBERT encodes each description $d_i$ using the BERT model to produce a dense vector representation $u_i$:
$$u_i = \text{BERT}(d_i)$$
3. **Siamese Network Structure**: SBERT uses a Siamese network structure, where two identical BERT networks encode a pair of sentences (descriptions) independently. The outputs are then used to compute similarity measures.
4. **Triplet Network Structure**: For training, SBERT uses triplet loss, which ensures that the embedding of a description is closer to its positive pair (similar description) than to a negative pair (dissimilar description) by a margin. The triplet loss $L$ is defined as:
$$L = \max(0, \|u_i - u_p\|^2 - \|u_i - u_n\|^2 + \alpha)$$
where $u_i$ is the anchor, $u_p$ is the positive example, $u_n$ is the negative example, and $\alpha$ is the margin.
5. **Cosine Similarity**: The resulting embeddings are compared using cosine similarity, which is given by:
$$\text{sim}(u_i, u_j) = \frac{u_i \cdot u_j}{\|u_i\|\|u_j\|}$$
where $u_i$ and $u_j$ are the embedding vectors of two descriptions.

### Comprehensive Analysis of Summarization Results

In addition to BART's performance on SPAGs 5-10, we also analyzed its performance on SPAGs 10-100 and compared it with the GPT-2 model. As shown in **Table S1**, BART consistently outperformed GPT-2 across both datasets, achieving a mean similarity score of 0.52 on SPAGs 10-100, while GPT-2 scored 0.37 and 0.33 on SPAGs 5-10 and SPAGs 10-100, respectively. The results indicate that BART, being a model tailored for summarization, is more effective in capturing essential information, particularly when the task involves summarizing larger node sets.

While these results demonstrate the potential of BART and GPT-2 for summarization tasks, it is important to note that this study primarily focuses on establishing a flexible and scalable framework for PAG summarization. We believe that the summarization performance can be significantly improved by replacing the current models with more advanced language models, such as Chat-GPT 4 or LLaMA 3. These models are better equipped to capture semantic nuances, which could further enhance the accuracy and coherence of the summaries. However, deploying these advanced models comes with significant computational requirements, which would need to be addressed in future implementations.

In conclusion, the current framework provides a robust foundation for summarizing biological data, and with the integration of more powerful language models, the system can deliver even more precise and informative summaries of biological processes.

| Evaluation Dataset | Models | Mean Similarity ± Std |
|---|---|---|
| SPAGs 5-10 | Bart | $0.63 \pm 0.16$ |
| | GPT2 | $0.37 \pm 0.16$ |
| SPAGs 10-100 | Bart | $0.52 \pm 0.18$ |
| | GPT2 | $0.33 \pm 0.17$ |

**Table S1.** Statistic Analysis for the summarization of two methods (BART and GPT-2) on different evaluation datasets SPAGs 5-10 and SPAGs 10-100)
